# Unearthing New Genomic Markers of Drug Response by Improved Measurement of Discriminative Power

**DOI:** 10.1101/033092

**Authors:** Cuong C. Dang, Antonio Peón, Pedro J. Ballester

**Author notes:** Email addresses: CCD, AP, PJB.

## Abstract

**Background:** Oncology drugs are only effective in a small proportion of cancer patients. Our current ability to identify these responsive patients before treatment is still poor in most cases. Thus, there is a pressing need to discover response markers for marketed and research oncology drugs in order to improve patient survival, reduce healthcare costs and enhance success rates in clinical trials. Screening these drugs against a large panel of cancer cell lines has been employed to discover new genomic markers of *in vitro* drug response, which can now be further evaluated on more accurate tumour models. However, while the identification of discriminative markers among thousands of candidate drug-gene associations in the data is error-prone, an appraisal of the effectiveness of such detection task is currently lacking.

**Results:** Here we present a new non-parametric method to measuring the discriminative power of a drug-gene association. This is enabled by the identification of an auxiliary threshold posing this task as a binary classification problem. Unlike parametric statistical tests, the adopted non-parametric test has the advantage of not making strong assumptions about the data distorting the identification of genomic markers. Furthermore, we introduce a new benchmark to further validate these markers *in vitro* using more recent data not used to identify the markers. The application of this new methodology has led to the identification of 128 new genomic markers distributed across 61% of the analysed drugs, including 5 drugs without previously known markers, which were missed by the MANOVA test initially applied to analyse data from the Genomics of Drug Sensitivity in Cancer consortium.

**Abbreviation:**

(WT): wild-type
(GDSC): Genomics of Drug Sensitivity in Cancer
(TP): true positive
(TN): true negative
(FP): false positive
(FN): false negative
(MCC): Matthews Correlation Co-efficient.

## Introduction

Cancer is a leading cause of morbidity and mortality in industrialised nations, with failed treatment being often life-threatening. While a wide range of drugs are now available to treat cancer patients, in practice only a small proportion of them respond to these drugs [1]. Worse yet, our current ability to identify responsive patients before treatment is still poor in most cases [2]. This situation has a negative impact on patient survival (the tumour keeps growing until an effective drug is administered), healthcare costs (very expensive drugs are ineffective, and thus wasted, on most cancer patients [1, 3]) and the success rates of oncology clinical trials (10% fall in Phase II studies, with the number of phase III terminations doubling in recent years [4]). Therefore, there is a pressing need to understand and predict this aspect of human variation to make therapy safer and more effective by determining which drugs will be more appropriate for any given patient.

The analysis of tumour and germline DNA has been investigated as a way to personalise cancer therapies for quite some time [5]. However, the recent and comprehensive flood of new data from much cheaper and faster Next Generation Sequencing technologies along with the maturity of more established molecular profiling technologies represents an unprecedented opportunity to study the molecular basis of drug response. These data have shown that drug targets often present genomic alterations across patient tumours [6]. At the molecular level, these somatic mutations affect the abundance and function of gene products driving tumour growth and hence may influence disease outcome and/or response to therapy [7]. Therefore, there is opportunity for genetic information to aid the selection of effective therapy by relating the molecular profile of tumours to their observed sensitivity to drugs. Research on the identification of drug-gene associations to be used as predictive biomarkers of *in vitro* drug response is carried out on human cancer tumour-derived cell lines [8–10]. Cell lines allow relatively quick and cheap experiments that are generally not feasible on more accurate disease models [11]. Here the molecular profile of the untreated cell line is determined and a phenotypic readout is measured to assess the intrinsic cell sensitivity or resistance to the tested drug. In addition to biomarker discovery [8–10], these data sets have also been used to enable pharmacogenomic modelling [12–14], pharmacotranscriptomic modelling [15, 16], QSAR modelling [17, 18], drug repositioning [18, 19] and molecular target identification [19–21], among other applications.

Our study focuses on the Genomics of Drug Sensitivity in Cancer (GDSC) data analysed by Garnett et al. [9] and publicly released after additional curation [22]. The released data set comprises 638 human tumour cell lines, representing a broad spectrum of common and rare cancer types. One benefit of looking at a large number of cell lines is that the pool of data becomes larger, which is crucial for *in vitro* biomarker discovery. These authors profiled each cell line for various genetic abnormalities, including point mutations, gene amplifications, gene deletions, microsatellite instability, frequently occurring DNA rearrangements and changes in gene expression. Next, the sensitivity of 130 drugs against these cell lines was measured with a cell viability assay *in vitro* (cell sensitivity to a drug was summarised by the half-maximal inhibitory concentration or IC_50_ of the drug-cell pair). A p-value was calculated for 8637 drug-gene associations using a MANOVA test (P_MANOVA_), with 396 of those associations being above a FDR=20% Benjamini-Hochberg [23] adjusted threshold and thus deemed significant (details in the Methods section). Overall, it was found that only few drugs had strong genomic markers, with no actionable mutations being identified for 14 drugs.

However, a statistically significant drug-gene association is not necessarily a useful genomic marker of *in vitro* drug response. There are two types of errors at this inter-association level: a false association (type I error or false positive) or a missed association (type II error or false negative). False negatives are the most worrying types of errors because these are hard to detect and can have particularly adverse consequences (e.g. missing a genomic marker able to identify tumours sensitive to a drug for which no marker has been found yet). Indeed, significant p-values are merely intended to highlight potential discoveries among thousands of possibilities and thus their practical importance still have to be evaluated for the problem at hand [24–26]. For example, a significant drug-gene association can be become non-significant with the availability of more data and hence be revealed as a spurious correlation. Another possibility is that the association is significant but its effect is tiny and thus of little consequence for identifying sensitive tumours. In this context, the practical importance of a potential marker is measured by how well the gene mutation discriminates between cell lines from an independent test set according their sensitivity to a given drug. Importantly, while a parametric test such as MANOVA makes strong modelling assumptions [27] (e.g. normality and equal variances in the distribution of residuals), the distribution of drug responses of the compared groups of cell lines is often skewed, contain outliers and/or have different variances. Consequently, p-values from the MANOVA test may be more prone to Type I and Type II errors than statistical tests requiring milder assumptions. Thus, research intended to identify more appropriate statistical procedures for biomarker discovery on comprehensive pharmacogenomic resources such as GDSC is crucial to make the most out of these valuable data.

Here we will investigate the impact that the choice of the statistical test has on the systematic identification of genomic markers of drug sensitivity on GDSC pharmacogenomic data. The assessment will be carried out by comparing drug-gene ssociations identified by the MANOVA test with those identified by Pearson’s chi-squared test. The latter is a non-parametric test [28] and hence it does not make strong modelling assumptions distorting the detection task. This chi-squared test is applied to binary classification and hence we propose here an auxiliary threshold to enable its application to this problem. Furthermore, the largest discrepancies between both statistical tests on the training data set will be visualised and discussed with respect to the discriminative power of its significant and non-significant drug-gene associations. In addition, we will introduce a benchmark using a more recent GDSC dataset than that employed for the identification of statistically significant drug-gene associations and use it to validate *in vitro* the single-gene markers arising from each statistical test. This is timely research because the issue of systematically validating markers *in vitro* has not been addressed yet and thus it is currently unknown to which extent the limitations of the statistical test affect genomic marker discovery.

## Results

### Improved measurement of discriminative power by the chi-squared test

Genomic markers of drug response aim at identifying gene alterations that best discriminate between tumours regarding their sensitivity to a given drug. The ANOVA family of statistical tests attempts to determine how discriminative is the gene alteration by comparing the intra-group variances with the inter-group variances based on several strong assumptions about the data [29]. In order to enable the application of the non-parametric chi-squared test, a suitable IC_50_ threshold is required to define two auxiliary classes of cell lines, those most sensitive to the drug and those most resistant to the drug, which permits posing biomarker evaluation as a binary classification problem. Such threshold cannot be a fixed IC_50_ value for all drugs due to the different IC_50_ ranges across drugs (otherwise, all cell lines would be sensitive to the most potent drugs). Likewise, it would not be meaningful to fix the same IC_50_ threshold for all drug-gene associations within a given drug (for example, using the mean of the drug’s IC_50_s along with a rare mutation would result in a threshold splitting the WT cell lines in about half regardless of how more sensitive the mutant cell lines could be). For each drug-gene association, this issue can be overcome by characterising the typical sensitivity of each group of cell lines (i.e. those with the mutated gene and those with the WT gene) and calculating the threshold as the mean of the sensitivities of both groups. However, if each group was characterised by the mean of its IC_50_ values, the presence of strong outliers and/or a highly skewed IC_50_ distribution would distort the position of the threshold. Thus, we characterise each group of cells by its median IC_50_ and define this mutation-dependent threshold as the mean of both medians (e.g. the dotted red line of the scatter plot in Figure 2A). This definition is advantageous in that the size of each group and their outliers do not alter the position of this decision boundary, which is equidistant to both classes and leads to an intuitive notion of class membership as distance from the threshold.

**Fig 1.**
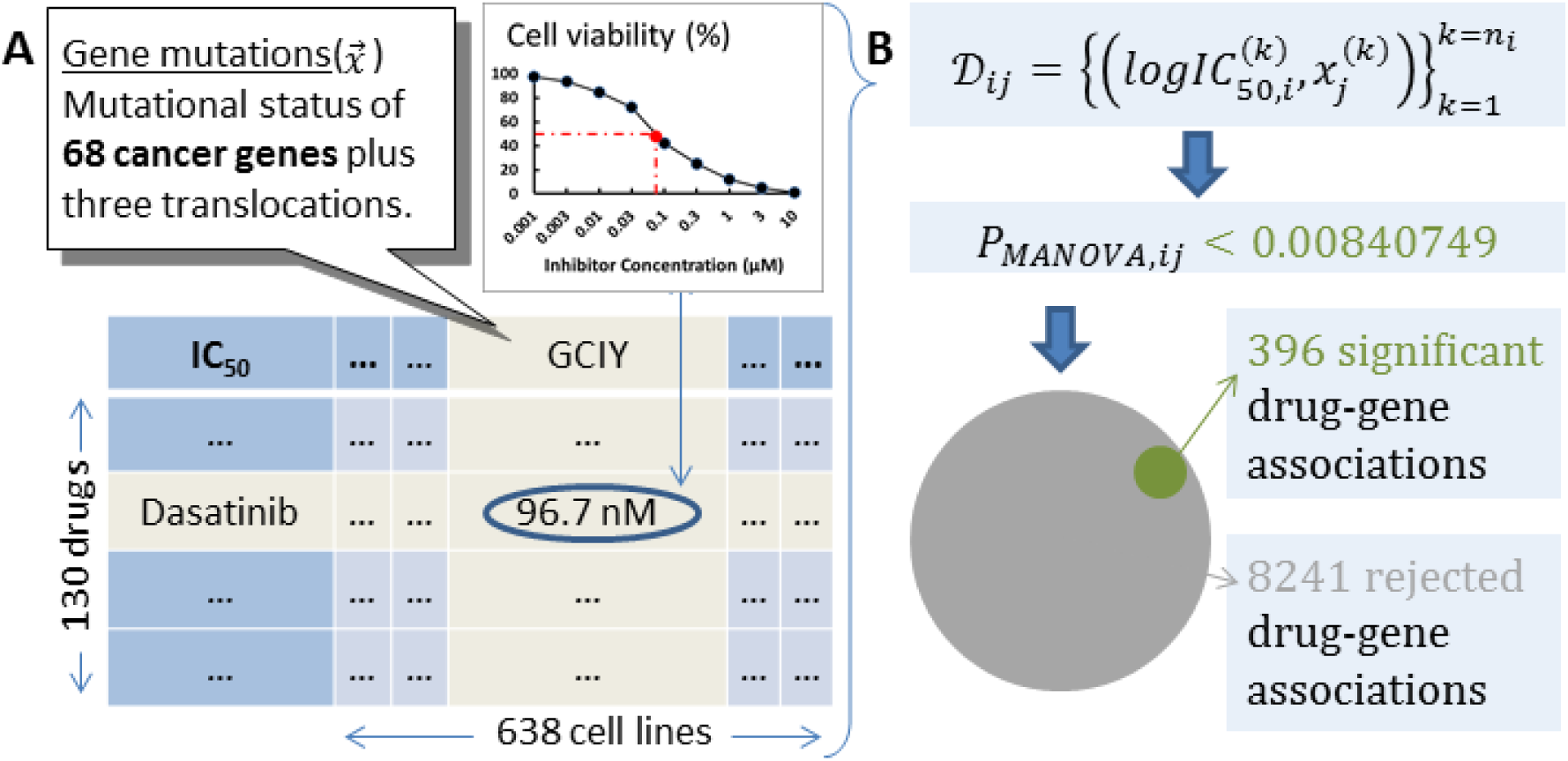
Released GDSC data. **(A)** Garnett et al.[9] analysed a slightly different dataset than the one that was later released. In the released dataset, a panel of 130 drugs was tested against 638 cancer cell lines, leading to 47748 IC_50_ values (57.6% of all possible drug-cell pairs). For each cell line, 68 cancer genes were sequenced and their mutational status determined, plus three translocations and a microsatellite instability status. **(B)** A dataset D_ij_ can be compiled for each drug-gene combination comprising the n_i_ cell responses to the i^th^ drug (in our case, each response as the logarithm base 10 of IC_50_ in μM units), with x_j_^(k)^ being a binary variable indicating whether the j^th^ gene is mutated or not in the k^th^ cell line. Next, a p-value was calculated for each drug-gene pair using the MANOVA test. Those pairs with p-values below an adjusted threshold of 0.00840749 were considered statistically significant (396 of the 8637 drug-gene associations).

**Fig 2.**
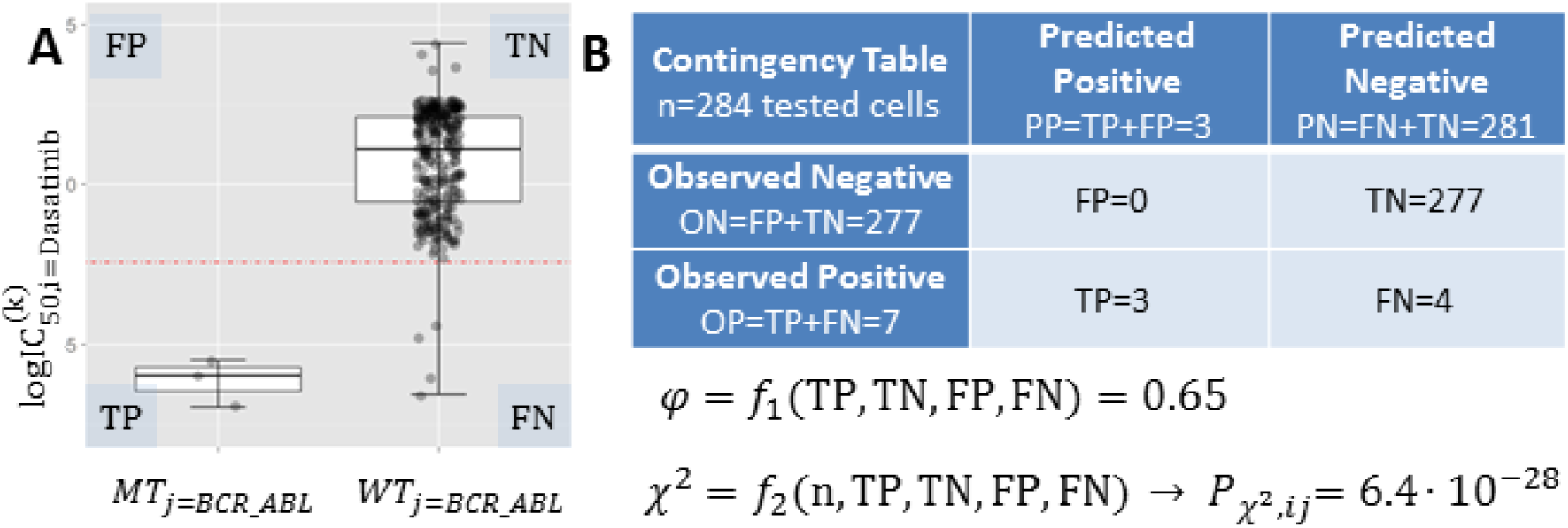
Measuring the discriminative power of a genomic marker with φ and the chi-squared test. **(A)** Scatter plot showing the logIC_50_ of n=284 cell lines screened against the marketed drug Dasatinib. The left boxplot shows BCR_ABL positive cell lines, whereas the boxplot on the right shows cell lines without this mutation (the median of each group appears as a black horizontal line within the boxplot). The red dotted line is the IC_50_ threshold, which is defined as the mean of both medians. **(B)** Contingency table showing the number of training set cell lines in each of the four non-overlapping categories (TP, FN, FP, TN), where positives are cell lines below the threshold in the case of a sensitising mutation (above the threshold if the mutation induces resistance). φ and 𝒳^2^ are functions of these metrics and summarise binary classification performance, as further described in the Methods section. BCR_ABL is a very strong marker of Dasatinib sensitivity as shown in the scatter plot and highlighted by both statistical tests (P_MANOVA_=1.4·10^-10^, P_*χ*^2^_=6.4·10^-28^), offering unusually high discrimination between cell lines according to their relative drug sensitivity (φ=0.65).

Once this IC_50_ threshold is calculated, the mutation-based prediction of drug response of a cell line can be categorised as a true positive (TP), true negative (TN), false positive (FP) or false negative (FN). These relative measures of drug sensitivity are only intended to quantify the discrimination between mutated and WT cell lines and must not be mistaken by absolute measures of drug sensitivity (e.g. a cell line can be defined as sensitive to a drug if its IC_50_ is better than the median IC_50_ of all cell lines for that drug, however such threshold may poorly measure how different are the drug responses of mutated and WT tumours). From this contingency table at the intra-association level, the discrimination offered by a drug-gene association can be summarised by its Matthews Correlation Coefficient (MCC) [30], as specified in the Methods section. Since cells are now partitioned into four non-overlapping categories with respect to their response to a drug, the chi-squared test statistic (denoted as 𝒳^2^) can be computed from this 2 × 2 contingency table to identify those drug-gene associations with statistically significant discriminative power (𝒳^2^ measures how far is the contingency table obtained by the classification method from the values that would be expected by chance). The process is sketched in Figure 2 and leads to an alternative set of p-values from the chi-squared test (P_*χ*^2^_), whose definitions and calculations are provided in the Methods section. To establish which associations are significant according to the chi-squared test, we also calculated for this case the FDR=20% Benjamini-Hochberg adjusted threshold (0.00940155). Overall, 403 statistically significant drug-gene associations were found using the chi-squared test from the same set of 8637 associations that were downloaded (i.e. seven significant associations more than with the MANOVA test). Importantly, only 171 associations of these markers were found by the MANOVA test. These deviations of the MANOVA test with respect to the results provided by the non-parametric test will be investigated in the next section to highlight potential false and missed biomarkers.

A last aspect to discuss about the proposed methodology is the duality of MCC and 𝒳^2^. In statistics, MCC is known as the φ coefficient, which was introduced [31] by Yule in 1912 and later rediscovered [30] by Matthews in 1975 as the MCC (interestingly, despite being more recent, the MCC has become a much more popular metric for binary classification than the φ coefficient [32–37]). As 𝒳^2^= n·φ^2^ holds[31], so does 𝒳^2^=n·MCC^2^ with n being the number of tested cell lines for the considered drug and thus MCC will be highly correlated with P_*χ*^2^_. To avoid confusion, we will use φ to refer to discrimination at this intra-association level (i.e. to identify the markers) and reserve MCC for the validation of the identified markers as a separate binary classification problem at the inter-association level that we will introduce later. Figure 3A presents the number of drug-gene associations for each number of tested cell lines, from which it is observed that each drug has only been tested on a subset n of the 638 cell lines (i.e. gene associations for a given drug will be all evaluated on the same number of cell lines n). Two distinctive groups of drugs emerge: those tested on around 300 cell lines (red bars) and those tested around 450 cell lines (black bars). Figure 3B shows that φ and-logP_*χ*^2^_ are highly correlated even across different n (for associations with the same n, a perfect Pearson and Spearman correlation is obtained as expected – data not shown). Given the observed φ distribution of n values, all markers with an φ of 0.15 or more are found unlikely to have arisen by chance. This connexion is useful in that φ is widely used [32–37] but without establishing its statistical significance for the tackled problem instance.

**Fig 3.**
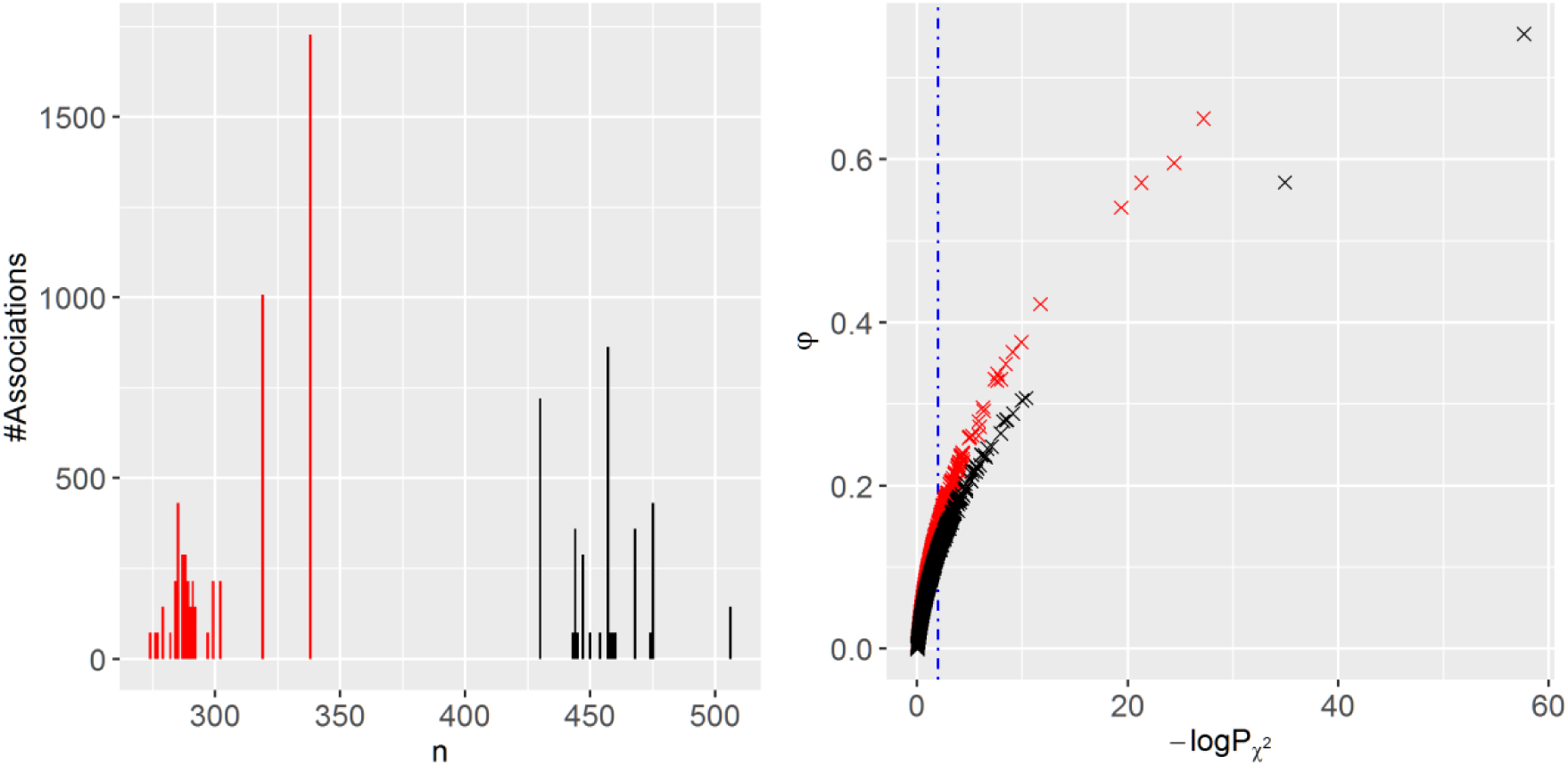
φ vs.-logP_*χ*^2^_ across all the 8637 drug-gene associations from GDSC. **(A)** Number of drug-gene associations for each number of tested cell lines (n). Two distinctive groups of drugs emerge: those tested on around 300 cell lines (red bars) and those tested around 450 cell lines (black bars). **(B)** φ versus-logP_*χ*^2^_ across the drug-gene associations (same colour code). The Spearman and Pearson correlations between both metrics are 0.99 and 0.82, respectively. The vertical blue line marks the significance cutoff for the chi-squared test. The plot shows that all markers with an φ of 0.15 or more are too discriminative to have arisen by chance (above an φ of 0.12 if we restrict to the markers evaluated with more data shown as black crosses).

### Potential false-positive and false-negative markers of the MANOVA test

We have introduced a new method directly measuring the discriminative power of a drug-gene association using the φ along with its significance using P_*χ*^2^_. We analyse next those associations where the MANOVA test deviates the most from this non-parametric test. First, we identified the association with the largest difference between P_MANOVA_ and P_*χ*^2^_ among those not significant by the chi-squared test. The left scatter plot in Figure 4 shows that this drug-gene association (GW441756-FLT3) discriminates poorly between mutant and WT cell lines despite a very low P_MANOVA_∼10^-10^. In contrast, a high P_*χ*^2^_∼10^-1^ is obtained which means that the chi-squared test rejected this potential false positive of the MANOVA test.

**Fig 4.**
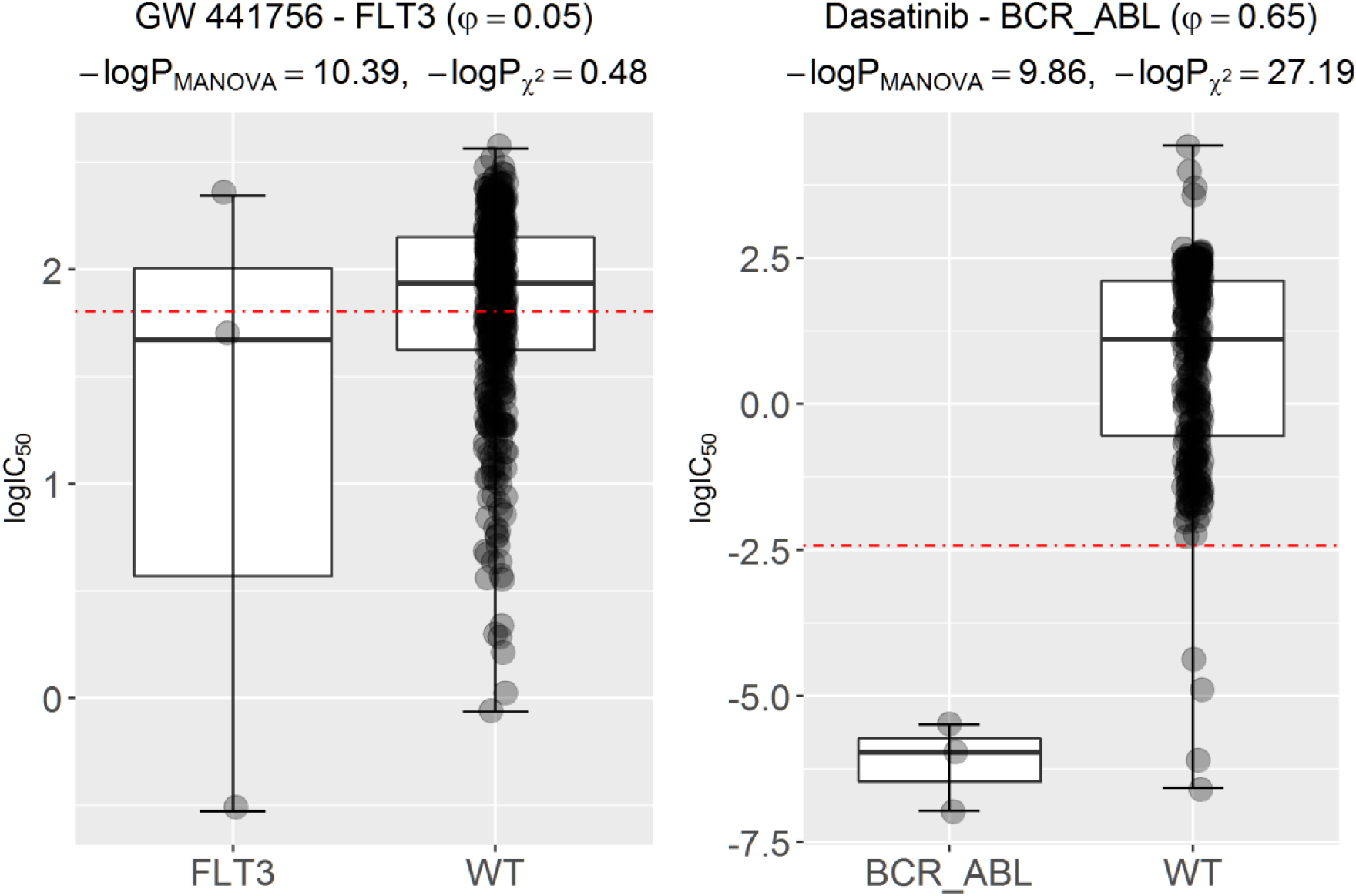
Potential false-positive marker of the MANOVA test incorrectly rejected by the chi-squared test. (**left**) The scatter plot for the drug-gene association (GW441756-FLT3) with the largest-logP_MANOVA_ among those not significant according to the chi-squared test. Hence, mutated-FLT3 is a marker of sensitivity to the experimental drug GW441756 according to the MANOVA test, but not according to the chi-squared test. In the plotted training set, this marker offers practically no discriminative power as further evidenced by a φ of just 0.05 and similar drug response (logIC_50_) distributions of mutated and WT cell lines. However, this marker provides an MCC of 0.10 on the test and hence this is a false negative of the chi-squared test. (**right**) Conversely, to assess the consistency of the MANOVA test, we searched for the drug-gene association with largest-logP_*χ*^2^_ among those with a similar-logP_MANOVA_ to that of GW441756-FLT3, which is Dasatinib-BCR_ABL. Whereas the p-value for Dasatinib-BCR_ABL is of the same magnitude as that for GW441756-FLT3 using the MANOVA test (P_MANOVA_∼10^-10^), the p-values for the same associations using the chi-squared test differ is almost 27 orders of magnitude. Thus, unlike the chi-squared test, the MANOVA test is unable to detect the extreme difference in discriminative power offered by these two drug-gene associations. Indeed, the BCR_ABL translocation is a highly discriminative marker of Dasatinib sensitivity (φ=0.65), as also evidenced by the barely overlapping drug response distributions from each set of cell lines. This is confirmed in the test set, where the Dasatinib-BCR_ABL marker obtains an MCC of 0.21.

Conversely, to assess the consistency of the MANOVA test, we searched for the drug-gene association with smallest P_*χ*^2^_ among those with a similar P_MANOVA_ to that of GW441756-FLT3, which is Dasatinib-BCR_ABL (Figure 4 right). The BCR_ABL translocation is a highly discriminative marker of Dasatinib sensitivity (φ =0.65), as evidenced by the barely overlapping drug response distributions from each set of cell lines. Note that, whereas the p-value for Dasatinib-BCR_ABL is of the same magnitude as that for GW441756-FLT3 using the MANOVA test (P_MANOVA_∼10^-10^), the p-values for the same associations using the chi-squared test are almost 27 orders of magnitude apart. Thus, unlike the chi-squared test, the MANOVA test is unable to detect the extreme difference in discriminative power offered by these two drug-gene associations.

The next experiment consists in searching for the largest discrepancy in the opposite direction. First, we identified the association with the largest difference between P_MANOVA_ and P_*χ*^2^_, this time among those not significant by the MANOVA test. The left scatter plot in Figure 5 shows marked difference in the two drug response distributions of this drug-gene association (Dasatinib-CDKN2a.p14), suggesting that this is a potential false negative of the MANOVA test despite a high P_MANOVA_∼10^-1^. In contrast, a low P_*χ*^2^_∼10^-9^ is obtained, which means that the chi-squared test detected this potential false negative of the MANOVA test. Conversely, to assess again the consistency of the MANOVA test, we searched for the drug-gene association with smallest P_MANOVA_ among those with a similar P_*χ*^2^_ to that of Dasatinib-CDKN2a.p14, which is SB590885-BRAF (Figure 5 right). Whereas the p-values for Dasatinib-CDKN2a.p14 and SB590885-BRAF differ 27 orders of magnitude using the MANOVA test, the p-values for the same associations have similar p-values using the chi-squared test P_*χ*^2^_∼^-9^. Thus, unlike the chi-squared test, the MANOVA test is unable to detect that both markers have similar discriminative power as also indicated by the MCC (SB590885-BRAF has a φ of 0.29 for 0.35 of Dasatinib-CDKN2a.p14).

**Fig 5.**
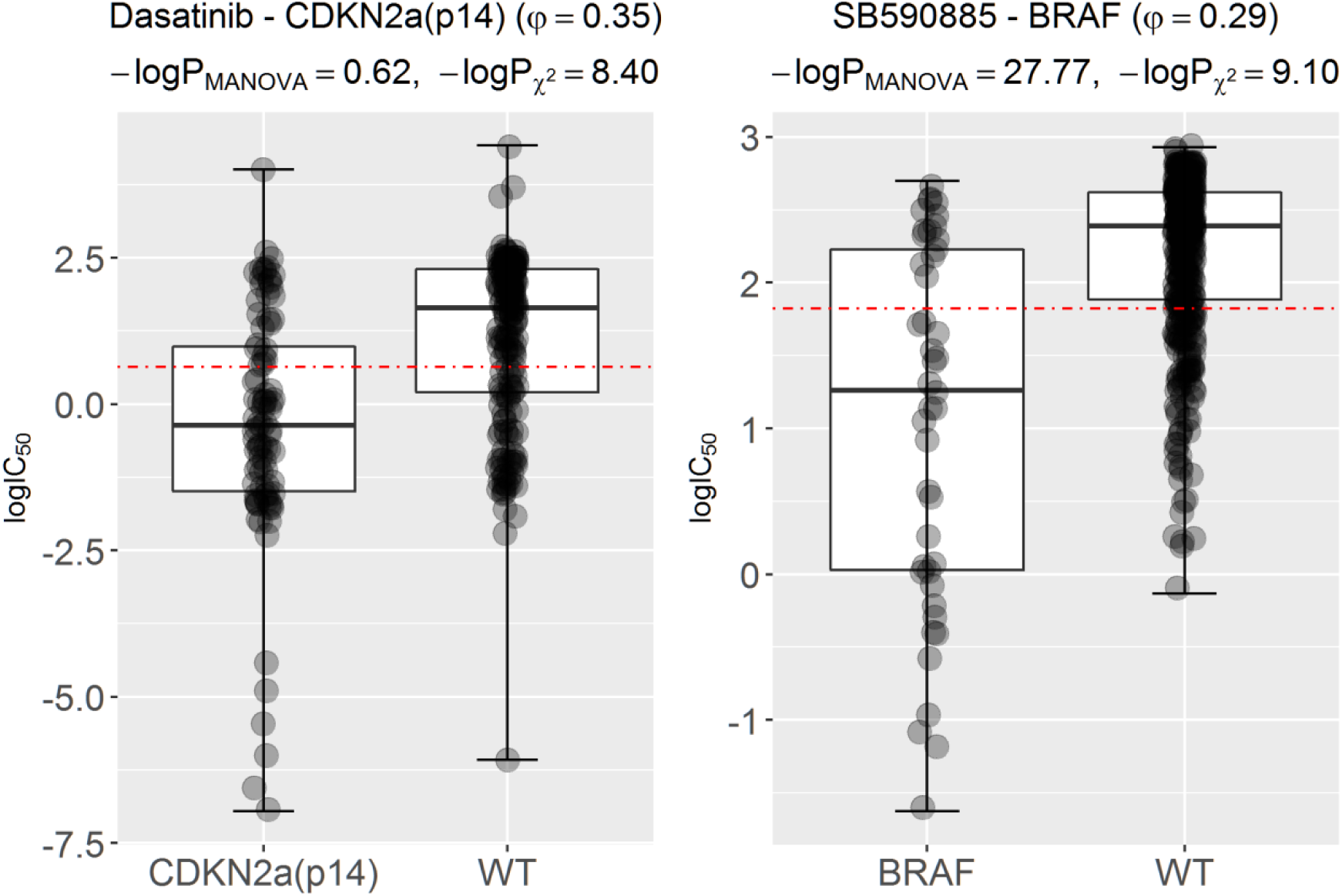
Potential false-negative marker of the MANOVA test detected by the chi-squared test. (**left**) The scatter plot for the drug-gene association (Dasatinib-CDKN2a.p14) with the largest-logP_*χ*^2^_ among those not significant according to the MANOVA test. Hence, mutated-CDKN2a.p14 is a potential marker of sensitivity to the marketed drug Dasatinib according to the chi-squared test, but not according to the MANOVA test. However, this marker has predictive value as it provides MCC=0.13 on the test set. Therefore, the chi-squared test detected this potential false negative of the MANOVA test. (**right**) Conversely, to assess the consistency of the MANOVA test, we searched for the drug-gene association with largest-logP_MANOVA_ among those with a similar P_*χ*^2^_ to that of Dasatinib-CDKN2a.p14, which is SB590885-BRAF. Whereas the p-values for Dasatinib-CDKN2a.p14 and SB590885-BRAF differ in 27 orders of magnitude using the MANOVA test, the p-values for the same associations have similar p-values using the chi-squared test (P_*χ*^2^_∽10^9^). Thus, unlike the chi-squared test, the MANOVA test is unable to detect that both markers have similar discriminative power (SB590885-BRAF has a φ of 0.29 for 0.35 of Dasatinib-CDKN2a.p14). SB590885-BRAF is a true positive of both tests as its MCC on the test set is 0.27.

### Validation of single-gene markers on a more recent GDSC data set

We propose a new benchmark based on using the most recent comparable GDSC data as test sets. For the 127 drugs in common between releases 1 and 5, two non-overlapping data sets are generated per drug. Training sets from data in release 1 along with their logIC_50_s for the considered drug, which were used to identify genomic markers as previously explained. Further, test sets contain the new cell lines tested with the drug in release 5. Thereafter, the significant drug-gene associations from each statistical test are evaluated on these test sets. A cell line sensitivity threshold was previously defined in order to discriminate between those resistant or sensitive to the considered drug. For each drug, we calculated the threshold as the median of all the logIC_50_ values from training set cell lines. Consequently, cell lines with logIC_50_ lower than such threshold are sensitive to the drug of interest, whereas those with logIC_50_ higher the threshold are resistant. Lastly, classification performance of a marker on its test set is summarised with the MCC.

Figure 6 presents a comparison between detection methods using this benchmark. The three compared methods are those based on the chi-squared test (B), the MANOVA test (C) and their consensus (A; the association is significant if it is significant by both tests). We can see that the consensus method is the most predictive (full results in additional file 1), followed by associations only significant with the chi-squared test (additional file 2) and those only significant by the MANOVA test (additional file 3). These results show that the overall predictive value of the markers revealed by the chi-squared test is higher than that arising from the MANOVA test and also that the consensus of both tests is more predictive than any of these two tests alone. While most of the markers provide better prediction than random classification (MCC=0 [38]), their generally low test set MCC values regardless of the employed detection method highlight how hard is to identify predictive markers of drug response.

**Fig 6.**
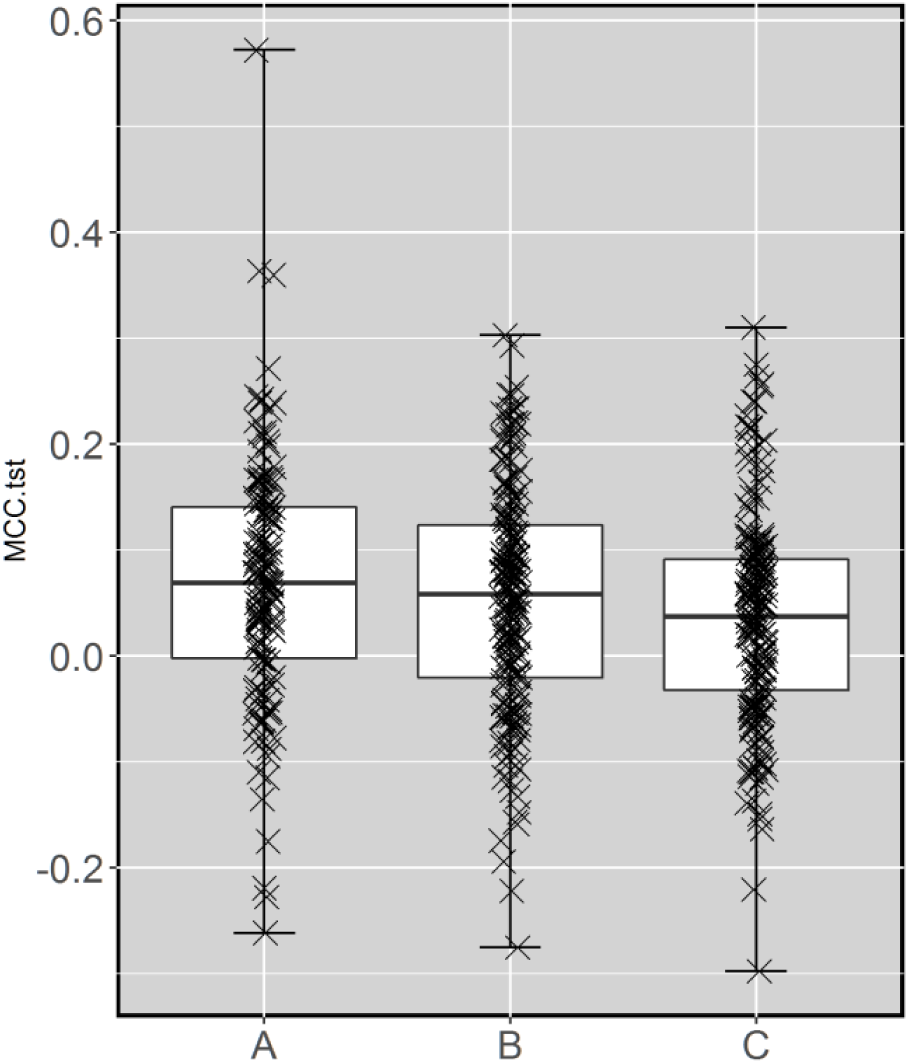
Test set performance of three methods to identify single-gene markers. The methods are evaluated by their ability to correctly classify more recently-tested cell lines as sensitive or resistant to the considered drug via the MCC on the test set. There is no overlap between test sets and those employed to identify all drug-gene associations (training sets). The three compared methods are those based on the chi-squared test (B), the MANOVA test (C) and their consensus (A; the association is significant if it is significant by both tests). We can see that the consensus method is the most predictive, followed by associations only significant with the chi-squared test (B) and those only significant by the MANOVA test (C). These results show that the overall predictive value of the markers revealed by the chi-squared test is higher than that arising from the MANOVA test and also that the consensus of both tests is more predictive than any of these two tests alone. While most of the markers provide better prediction than random classification (MCC=0), their generally low test set MCC values regardless of the employed detection method highlight how hard is to identify predictive markers of drug sensitivity.

We also use this framework to further validate *in vitro* the markers shown in Figures 4 and 5 as examples. The GW441756-FLT3 marker provides an MCC of 0.10 on the test despite having weak discriminative power on the training set and hence this is a false negative of the chi-squared test. The Dasatinib-BCR_ABL marker obtains an MCC of 0.21 on the test set. Dasatinib-CDKN2a.p14 provides MCC=0.13 on the test set. Therefore, the chi-squared test detected this confirmed false negative of the MANOVA test. SB590885-BRAF is a true positive of both tests since its MCC on the test set is 0.27.

### 128 new markers unearthed by the chi-squared test and validated *in vitro*

The rest of the study will focus on unearthing these missed discoveries using the chi-squared test and further *in vitro* validation based on a test set made of more recent GDSC data. Indeed, these new genomic markers constitute additional knowledge that can be extracted from existing data, i.e. without requiring any further experiment. In the data released by the GDSC, the 396 genomic markers from the MANOVA test were distributed among 116 drugs, leaving the remaining 14 drugs without any maker. Of the 403 single-gene markers identified by the chi-squared test, 187 were not found by the MANOVA test and could not be evaluated on the test set because there are only 127 drugs in common between the training and test sets and some markers did not have mutant test set cell lines (i.e. test set MCC cannot be evaluated for these markers because these yield no prediction). For the same reasons, the situation is similar for the MANOVA test: only 182 of the 396 MANOVA-significant drug-gene associations were not found by chi-squared test and could not be evaluated on the test set. Further, there are 128 of the 187 associations from the chi-squared test with test set MCC greater than zero (115 of the 182 associations from the MANOVA test).

Figure 7 shows two examples of new chi-squared markers for drugs with previously-proposed MANOVA markers. The scatter plot at the top left identified the mutational status of CDK2NA as a new marker of sensitivity to Temsirolimus, which was missed by the MANOVA test. This marker predicts well which cell lines are sensitive to this drug (MCC of 0.30 on the test set; top right plot). The second example is shown at the bottom of Figure 7. The EWS_FLI1 translocation is also a new response marker for the drug BMS-754807, which was also missed by the MANOVA test. This marker provides good predictive performance on cell lines not used to identify the markers (MCC of 0.25 on the test set; bottom right plot). Overall, we have found new markers unearthed by the chi-squared test in 77 of the 127 drugs (see additional file 2).

**Fig 7.**
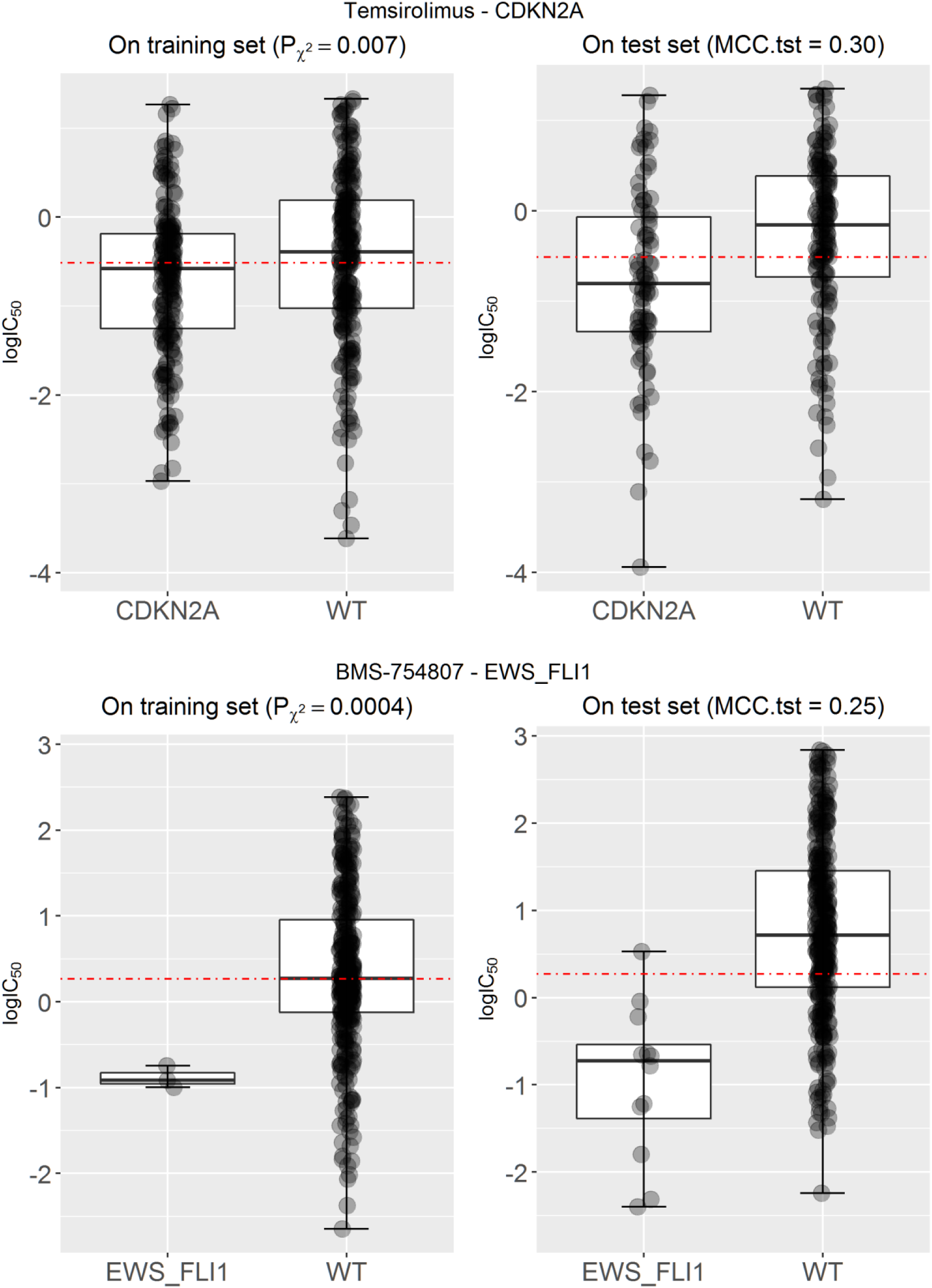
Examples of new genomic markers for drugs with previously-proposedMANOVA markers. **(top)** The mutational status of the CDKN2A gene is found to be the most discriminative marker for the approved drug Temsirolimus (MCC=0.30 on the test set,), which was missed by the MANOVA test (P_MANOVA_=9·10^-3^). **(bottom)** The EWS_FLI1 translocation is found to be the most discriminative marker for the development drug BMS-754807 (MCC=0.25 on the test set), which was also missed by the MANOVA test (P_MANOVA_=0.01). While both tests are being applied to exactly the same data, only the chi-squared test could identify these confirmed false negatives of the MANOVA test.

New genomic markers are particularly valuable in those drugs for which no marker is known yet. From our analysis, we have also identified seven new markers with MCC better than random classification for the five drugs for which the MANOVA test did not find any potential marker [9]: NU-7441, Cyclopamine, BI-2536, Gemcitabine and Epothilone B (see Additional file 2). Figure 8 shows the performance of two of these markers. On the right, the mutational status of the NOTCH1 gene is the most discriminative marker for the development drug BI-2536 (MCC=0.23 on the test set). On the left, EWS_FLI1-positive cell lines exhibit increased sensitivity to Gemcitabine (MCC=0.18 on the test set).

**Fig 8.**
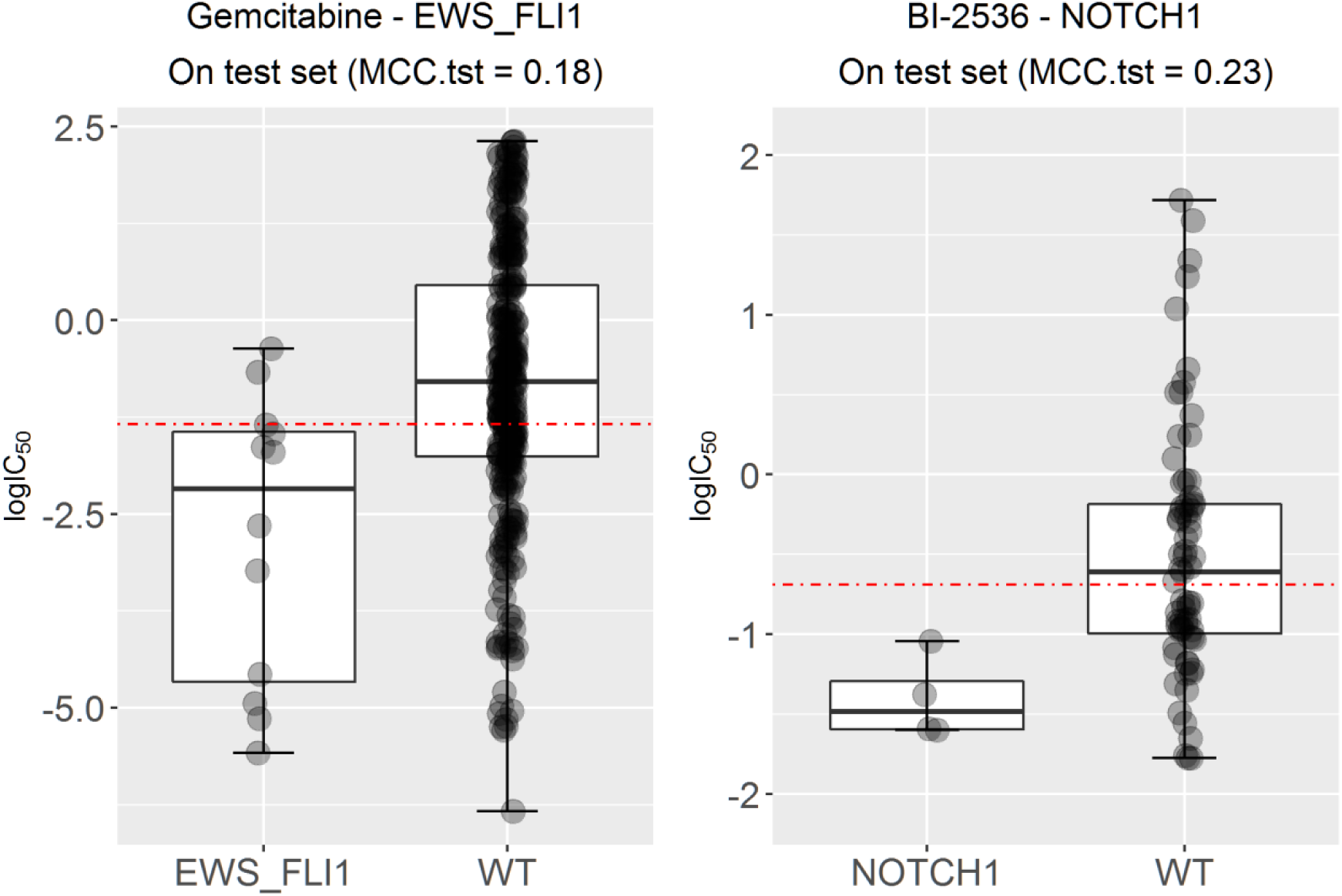
Examples of new genomic markers for drugs without previously-proposed known MANOVA markers. **(left)** The EWS_FLI1 translocation is found to be the most discriminative marker for the approved drug Gemcitabine (MCC=0.18 on the test set), which was missed by the MANOVA test (P_MANOVA_=0.06). **(right)** The mutational status of the NOTCH1 gene is found to be the most discriminative marker for the development drug BI-2536 (MCC=0.23 on the test set), which was also missed by the MANOVA test (P_MANOVA_=0.03).

## Discussion

To improve the search of genomic markers of drug response, we have presented a new non-parametric approach that directly measures the discriminative power of a drug-gene association by posing it as a binary classification problem. This change of perspective has been enabled by the introduction of an auxiliary threshold that is tailored to each association. Thus, discrimination can be measured with the 𝒳^2^ statistic and its significance with the chi-squared test, which provides a better alignment of the statistical and biological significance of a drug-gene association. Furthermore, we have shown that, since φ is linked to 𝒳^2^, the significance of a φ value can also be calculated with the chi-squared test.

Next, the chi-squared test has been applied to the identification of genomic markers from GDSC data and these markers compared to those arising from the MANOVA test[9]. Unlike the chi-squared test, statistical tests from the ANOVA family are parametric and thus expected to lead to inaccuracies when the data do not conform to the underlying modelling assumptions [27, 28]. Unlike the MANOVA test, the chi-squared test has the drawback of requiring the binarisation of logIC_50_ values, which leads to all misclassification errors having the same weight on the chi-squared test statistic regardless of the magnitude of this error. The largest discrepancies arising from both sets of p-values have been discussed in detail as shown in Figures 4 and 5, which provide examples of false negatives of both tests. False positive markers of either test are less important because they do not represent new knowledge, but resource-consuming false alarms, and may also become true positives with the arrival of more data.

Using the new benchmark, we have carried out a systematic comparison across 8637 drug-gene associations for which a p-value from the MANOVA test had been calculated in the GDSC study[9]. The MANOVA test highlighted 396 of these associations as statistically significant, for 403 from the chi-squared test looking at the same data. However, only 171 associations were deemed statistically significant by both tests. Ultimately, we have found that 216 of the 396 MANOVA-significant markers offer better than random performance. These drug-gene associations are those with positive MCC in additional files 1 and 3.

We have also found that 229 of the 403 𝒳^2^-significant markers offer better than random performance. Of these 229, 128 are new markers only detected by the chi-squared test (see additional file 2) and hence are false negatives of the MANOVA test. Temsirolimus-CDK2NA, 17AAG-CDK2NA or BMS-754807-EWS_FLI1 are among the most predictive of these new *in vitro* markers. Furthermore, we also identified 7 new markers with MCC better than random classification for the 5 drugs for which the MANOVA test did not find any marker [9]: NU-7441, Cyclopamine, BI-2536, Gemcitabine and Epothilone B. Overall, the predictive value of the markers revealed by the chi-squared test is higher than that arising from the MANOVA test and also that the consensus of both tests is more predictive than any of these two tests alone (see Figure 6). The former means that the chi-squared test should be preferred over the MANOVA test for this problem, the latter showing that the consensus of both tests highlights markers that are more likely to be predictive than those that are significant by only one of the tests.

Regarding best practices to compare two statistical tests for biomarker discovery, it could be argued that it is better to base the comparison on the ability of the tests to identify clinical markers. However, there are several reasons why this is inadequate. First of all, only a fraction of GDSC drugs have FDA-approved markers. Second, whereas clinical markers are so discriminative that are easily found by both methods, the challenge is to identify more subtle markers in the data. Indeed, the goal of the GDSC study was to search for still unknown markers to increase the ratio of patients that could benefit from personalised treatments (low for most clinical markers) as well as to find new markers for those drugs without clinical markers. Lastly, a gene mutation discriminative of *in vitro* drug response may be discriminative of human drug response, without still having been assessed in the clinic. A validation based on comparing the tests on clinical markers will be thus blind to the MANOVA test missing these discoveries.

Predictive biomarkers are highly sought after in drug development and clinical research [39, 40]. A vast amount of cancer genomics data is nowadays being generated [41] and thus there is an urgent need for their accurate analysis [42]. In the area of drug sensitivity marker discovery, recent multilateral efforts have been made [43, 44] to investigate the consistency of high-throughput pharmacogenomic data, which are collectively important to promote an optimal use of this valuable data by the relevant communities [45]. However, the impact of the strong modelling assumptions made by standard parametric tests on the discovery of genomic markers from data has not been analysed until now. Therefore, this study is important in a number of ways. First, these new genomic markers of *in vitro* drug response represent testable hypothesis that can now be evaluated on more relevant disease models to humans. Second, they may also constitute further evidence supporting newly proposed oncology targets [46]. Third, beyond the exploitation of these results, the widespread application of this methodology should lead to the discovery of new predictive biomarkers of *in vitro* drug response on existing data, as it has been the case here with the GDSC. Indeed, this new approach has been demonstrated on a large-scale drug screening against human cancer cell lines, but it can also be applied to other biomarker discovery problems such as those adopting more accurate disease models (e.g. primary tumours [47, 48], patient-derived xenografts [49, 50] or patients [51, 52]), those employing other molecular profiling data (e.g. transcriptomics [53], secretome proteomics [54], epigenomics [55] or single-cell genomics [56]) or those involving drug combinations [57]. Looking more broadly, the methodology can also be applied to large-scale drug screening against human or non-human molecularly-profiled pathogen cultures, such as those in antibacterial or agricultural research.

## Methods

### GDSC data

From release 1.0 of the Genomics of Drug Sensitivity in Cancer (GDSC) [22], we downloaded the following data files: gdsc_manova_input_w1.csv and gdsc_manova_output_w1.csv.

In gdsc_manova_input_w1.csv, there are 130 unique drugs as camptothecin was tested twice, drug ids 195 and 1003, and thus we only kept the instance that was more broadly tested (i.e. drug ID 1003 on 430 cell lines). Thus, effectively a panel of 130 drugs was screened against 638 cancer cell lines, leading to 47748 IC_50_ values (57.6% of all possible drug-cell pairs). Downloaded “IC_50_” values are more precisely the natural logarithm of IC_50_ in μM units (i.e. negative values represent drug responses more potent than 1μM). We converted each of these values into their logarithm base 10 in μM units, which we denote as logIC_50_ (e.g. logIC_50_=1 corresponds to IC_50_=10μM), as in this way differences between two drug response values are directly given as orders of magnitude in the molar scale.

gdsc_manova_input_w1.csv also contains genetic mutation data for 68 cancer genes, which were selected as the most frequently mutated cancer genes [9], characterising each of the 638 cell lines. For each gene-cell pair, a ‘x::y’description was provided by the GDSC, where ‘x’ identifies a coding variant and ‘y’ indicates copy number information from SNP6.0 data. As in Garnett et al. [9], a gene for which a mutation is not detected is considered to be wild-type (wt). A gene mutation is annotated if: a) a protein sequence variant is detected (x ≠{wt,na}) or b) a deletion/amplification is detected. The latter corresponds to a copy number (cn) variation different from the wt value of y=0<cn<8. Furthermore, three translocations were considered (BCR_ABL, MLL_AFF1 and EWS_FLI1). For each of these gene fusions, cell lines are identified as fusion not-detected or the identified fusion is given (i.e. wt or mutated with respect to the gene fusion, respectively). The microsatellite instability (msi) status of each cell line was also determined. Full details can be found in the original publication [9].

### Statistically significant drug-gene associations with the MANOVA test

Garnett *et al.*[9] carried out a fixed-effects MANOVA statistical test based on the genomic features specified in the previous section. An nx2 dose–response matrix consisting of IC_50_ and slope parameter for the n cell lines was constructed for each drug. A linear (no interaction terms) model was claimed to explain these observables from the genomic features as input and the tissue type as co-variate. Since this procedure was not fully specified (e.g. no test statistic choice or implementation information was provided), we used their results (gdsc_manova_output_w1.csv) and hence we did not recalculate them. This file contains 8701 drug-gene associations with p-values. As we are considering all those involving the 130 unique drugs (i.e. removing the camptothecin duplicate), we are left with 8637 drug-gene associations with p-values of which 396 were above a FDR=20% Benjamini-Hochberg adjusted threshold (0.00840749) and thus deemed significant according to this test. As usual [9], each statistically significant drug-gene association was considered to be a genomic marker of *in vitro* drug response.

### Measuring the discriminative power of a genomic marker with the chi-squared test

Let the training data for the association between the i^th^ drug and the j^th^ gene be

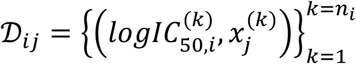

where *n*_*i*_ is the number of cell lines screened against the i^th^ drug and *k* denotes the considered cell line. The sets of mutated and WT cell lines with respect to the j^th^ gene, MT_j_ and WT_j_, be

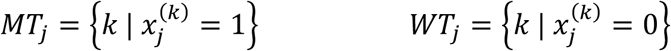

Next, the logIC_50_ threshold is defined as the average of the two median responses from each set (see subsection “Improved measurement of discriminative power by the chi-squared test”).

Thus, for each association between the i^th^ drug and the j^th^ gene, two steps are carried out to pose its evaluation as an intra-association binary classification problem.

Step 1: 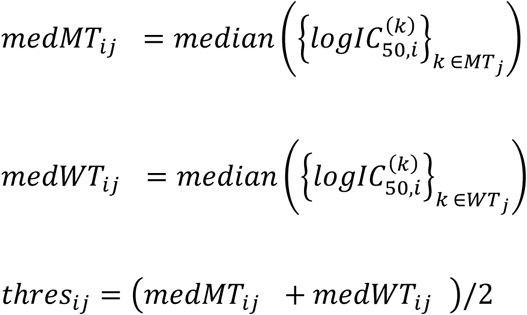
Step 2: if (*medMT*_*ij*_ < *medWT*_*ij*_) then mutant cell lines tend to be more sensitive to the drug and hence this is a genomic marker of drug sensitivity. Consequently, positives are defined as cell lines with logIC_50_< *thres*_*ij*_ and negatives are defined as cell lines with logIC_50_ ≥ *thres*_*ij*_. else if (*medMT*_*ij*_ ≥ *medWT*_*ij*_) then mutant cell lines tend to be more resistant to the drug and hence this is a genomic marker of drug resistance. Therefore, negatives are defined as cell lines with logIC_50_ < *thres*_*ij*_ and positives are defined as cell lines with logIC_50_ ≥ *thres*_*ij*_.

At this point, the set of all the cell lines tested with a given drug can be partitioned into four categories as defined in Figure 2: true positive (TP), true negative (TN), false positive (FP) or false negative (FN). From this contingency table, the discrimination offered by a drug-gene association can be summarised by the Matthews Correlation Coefficient (MCC) [30]

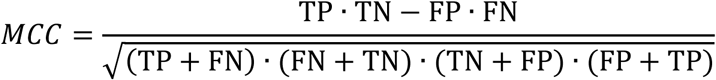

By the above definition of positives and negatives, MCC can only take values from 0 (gene mutation has absolutely no discriminative power) to 1 (gene mutation perfectly predicts whether cell lines are sensitive or resistant to the drug). Also, note that both the definition of the logIC_50_ threshold and the existence of mutated and wt cell lines in every association guarantees a non-zero value of the denominator in the MCC formula and thus MCC is always defined in this study. As previously explained, we report MCC as φ whenever this is calculated with the mutation-dependent threshold on training data (i.e. GDSC release 1.0).

### Statistically significant drug-gene associations with the chi-squared test

For each of the 8637 drug-gene associations, the chi-squared test statistic was computed from the 2x2 contingency table [29] to identify those drug-gene associations with statistically significant discriminative power. The formula to compute the chi-squared test statistic is

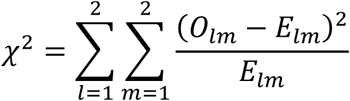

where O_lm_ are the four categories in the table (TP,TN,FN,FP) and E_Im_ are the corresponding expected values under the null hypothesis that this partition has arisen by chance. Thus, expected values are calculated with

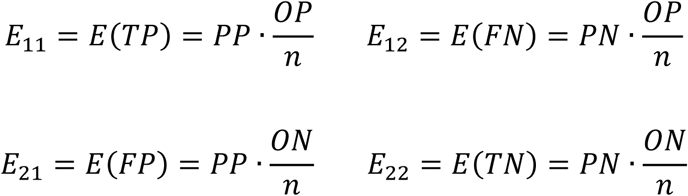

For instance, the expected value of TP, E(TP), is the number of predicted positives (PP) times the probability of a cell being a positive given as the proportion of observed positives (OP) in the n tested cells.

This chi-squared test statistic follows a 𝒳^2^ distribution with one degree of freedom and thus each p-value was computed with the R package *pchisq* from its corresponding 𝒳^2^ value, 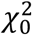 as

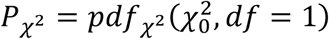

where *pdf*_*χ*^2^_ is the probability density function of the chi-square distribution. The process is sketched in Figure 2 and leads to an alternative set of p-values from the chi-squared test (P_*χ*^2^_). To establish which associations are significant according to the chi-squared test, we also calculated for this case the FDR=20% Benjamini-Hochberg adjusted threshold (0.00940155), that is

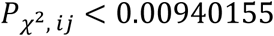

To facilitate reproducibility and the use of this methodology to analyse other pharmacogenomics data sets, the R script to calculate φ, chi-squared test statistic and P_*χ*^2^_ from gdsc_manova_input_w1.csv is available on request.

### Benchmark to validate genomic markers on more recent GDSC data

This benchmark is based on using more recent GDSC data as test sets. With this purpose, we downloaded new data from the latest release using the same experimental techniques to generate pharmacogenomic data and panel of selected genes as in release 1 (gdsc_manova_input_w5.csv). This release 5 contains 139 drugs tested on 708 cell lines comprising 79,401 logIC_50_ values (80.7% of all possible drug-cell pairs). For the 127 drugs in common between releases 1 and 5, two non-overlapping data sets are generated per drug. Training sets using data in release 1 (the minimum, average and maximum numbers of cell lines across training data sets are 237, 330 and 467, respectively), along with their logIC_50_s for the considered drug. These sets were used to identify genomic markers as previously explained. Test sets contain the new cell lines tested with the drug in release 5 (the minimum, average and maximum numbers of cell lines in the test data sets are 42, 171 and 306, respectively). Thus, a total of 254 data sets were assembled and analysed for this study.

The significant drug-gene associations from each statistical test are next evaluated on these test sets (this is the inter-association classification problem). A cell line sensitivity threshold was previously defined to discriminate between those resistant or sensitive to a given drug. For each drug, we calculated the threshold as the median of all the logIC_50_ values from training set cell lines. Consequently, cell lines with logIC_50_ lower than such threshold are sensitive to the drug of interest, whereas those with logIC_50_ higher the threshold are resistant. Lastly, classification performance of a marker on its test set is summarised with the MCC (this is different from φ, which has the same expression but uses a different threshold aimed instead at measuring the degree of separation between mutant and WT cell lines in the training set).

## Competing interests

The authors declare that they have no competing interests.

## Availability of data and materials

Data analysed in this paper was downloaded from releases 1.0 and 5.0 of the GDSC (ftp://ftp.sanger.ac.uk/pub4/cancerrxgene/releases/). All the results are compiled in the three additional files accompanying this paper.

## Ethics and consent statement

Not applicable.

## Author contributions

P.J.B. conceived the study, designed its implementation and wrote the manuscript. C.C.D. implemented the software and carried out the numerical experiments with the assistance of A.P. All authors discussed results and commented on the manuscript.

## Acknowledgements

We thank Gustavo Stolovitzky (IBM Research, NY, USA) for early feedback on the study. This work has been carried out thanks to the support of the A*MIDEX grant (n° ANR-11-IDEX-0001-02) funded by the French Government "Investissements d’Avenir» programme.

## Additional files

**Additional file 1 – results.127drugs.A-Consensus.xls**

Number of training cell lines (nTrain), prevalence of gene mutation, p-values, number of test set cell lines (nTest) and MCC on the test set (MCC.tst) for each significant drug-gene association from the consensus method evaluated on the test set.

**Additional file 2 – results.127drugs.B-ChiSquare.xls**

Number of training cell lines (nTrain), prevalence of gene mutation, p-values, number of test set cell lines (nTest) and MCC on the test set (MCC.tst) for each significant drug-gene association from the chi-squared test evaluated on the test set.

**Additional file 3 – results.127drugs.C-MANOVA.xls**

Number of training cell lines (nTrain), prevalence of gene mutation, p-values, number of test set cell lines (nTest) and MCC on the test set (MCC.tst) for each significant drug-gene association from the MANOVA test evaluated on the test set.

